# Ultralarge Modulation of Single Wall Carbon Nanotube Fluorescence Mediated by Neuromodulators Adsorbed on Arrays of Oligonucleotide Rings

**DOI:** 10.1101/351627

**Authors:** Abraham G. Beyene, Ali A. Alizadehmojarad, Gabriel Dorlhiac, Aaron M. Streets, Petr Král, Lela Vuković, Markita P. Landry

**Affiliations:** Department of Chemical and Biomolecular Engineering, University of California, Berkeley, Berkeley, CA 94720; Department of Chemistry and Biochemistry, University of Texas at El Paso, El Paso, TX 79968; Berkeley Biophysics Program, University of California, Berkeley, Berkeley, CA 94720; Department of Bioengineering, University of California, Berkeley, Berkeley, CA 94720; Chan-Zuckerberg Biohub, San Francisco, CA 94158; Department of Chemistry, Physics, and Biopharmaceutical Sciences, University of Illinois at Chicago, Chicago, IL 79968; California Institute for Quantitative Biosciences (qb3), University of California, Berkeley, Berkeley, CA 94720

## Abstract

Non-covalent interactions between single-stranded DNA (ssDNA) oligonucleotides and single wall carbon nanotubes (SWNTs) have provided a unique class of tunable chemistries for a variety of applications. However, mechanistic insight into both the photophysical and intermolecular phenomena underlying their utility is lacking, resulting in obligate heuristic approaches for producing ssDNA-SWNT based technologies. In this work, we present an ultrasensitive “turn-on” nanosensor for neuromodulators dopamine and norepinephrine with strong ΔF/F_0_ of up to 3500%, a signal appropriate for *in vivo* imaging, and uncover the photophysical principles and intermolecular interactions that govern the molecular recognition and fluorescence modulation of this nanosensor synthesized from the non-covalent conjugation of (GT)_6_ ssDNA strands on SWNTs. The fluorescence modulation of the ssDNA-SWNT conjugate is shown to exhibit remarkable sensitivity to the ssDNA sequence chemistry, length, and surface density, providing a wealth of parameters with which to tune nanosensor dynamic range and strength of fluorescence turn-on. We employ classical and quantum mechanical molecular dynamics simulations to rationalize our experimental findings. Calculations show that (GT)_6_ ssDNA form ordered loops around SWNT, inducing periodic surface potentials that modulate exciton recombination lifetimes. Further evidence is presented to elucidate how analyte binding modulates SWNT fluorescence. We discuss the implications of our findings for SWNT-based molecular sensing applications.

## Main

Single wall carbon nanotubes exhibit advantageous electronic and photophysical properties that make them attractive for a diverse field of applications in electronics^1–5^, sensing^6–9^, imaging^10–12,13^,and molecular transport^14–16^. SWNT fluorescence originates from radiative recombination of one-dimensional confined excitons, exhibits exceptional photostability, and is remarkably sensitive to the nanotube geometric and electronic structure as well as the local chemical environment.^17–19^ The sensitivity of SWNT fluorescence to the local chemical environment has been leveraged for the synthesis of optical probes in which polymer functionalizations serve a dual purpose of forming stable SWNT colloidal suspensions and conferring selective molecular recognition capabilities.^9,20^ Several SWNT-based probes with selective analyte mediated modulations in fluorescence quantum yield or shifts in optical band gaps on the order of ΔF/F_0_ between 9% and 80% have been reported.^9,21–25^

For *in vivo* molecular sensing applications, synthesizing suitable elements capable of transducing *in vivo* signals constitutes a formidable challenge.^26^ The spatiotemporal sensitivity required for *in vivo* utility – in particular for fast processes such as chemical neurotransmission in the brain – must account not just for analyte concentration levels, but also for the spatial spread of the signal (micrometers) as well as its temporal duration (milliseconds).^27,28^ An ideal probe therefore must satisfy several requirements, including high sensitivity, molecular selectivity, and optimal binding kinetics among others. The versatility and ease with which SWNTs can be functionalized by a wide range of polymers provides a great opportunity for a rational design of synthetic optical probes capable of detecting biomolecules in their native environment.^8^ However, despite proliferating reports of SWNT-polymer conjugates for biomolecule sensing, a robust pathway for translating SWNT nanosensors into *in vivo* sensing applications remains elusive. We identify two specific limitations in the development of SWNT based optical probes – lack of a rational design principle and dearth of *in vivo* implementation – and posit that a lack in fundamental understanding of how SWNT-polymer hybrid nanomaterials undergo selective fluorescence modulation by molecular targets underlies this limitation. This knowledge gap is evident in the status quo for nanosensor discovery, which relies on low-throughput screening techniques, and an inability to tune nanosensor performance once a discovery has been made.

In this work, we report a high turn-on nanosensor for neuromodulators dopamine and norepinephrine. We demonstrate how we can tune SWNT exciton recombination rates to increase nanosensor analyte sensitivity by over an order of magnitude compared to a previously reported nanosensor.^21^ Sequence-specific ‘short’ ssDNA polymers produced strongly quenched SWNT baseline fluorescence and a robust turn-on response to neuromodulators dopamine and norepinephrine. We find this phenomenon to be sensitive to the base sequence chemistry, polymer contour length, and surface density. Classical molecular dynamics (MD) calculations identified polymer-induced ‘electrostatic footprinting’ on the SWNT surface that induce periodic charge density isosurfaces. The surface potentials modulate exciton recombination and play a critical role in setting the baseline fluorescence of the ssDNA-SWNT conjugate. Further experimental and quantum mechanical MD (QMMD) simulations suggest a mechanism by which dopamine causes recovery of SWNT fluorescence. Experiments revealed the presence of specific molecular recognition sites in the ssDNA-SWNT corona that stabilize the surface adsorbed polymer when occupied by dopamine and norepinephrine analytes. QMMD simulations show that adsorbed dopamine analytes perturb the periodicity of the polymer induced SWNT surface potentials, allowing a competitive radiative relaxation of excitons and a fluorescence turn-on.

## Strong Fluorescent “Turn-on” Neuromodulator Nanosensors

Prior work has shown the fluorescence intensity of (GT)_15_-SWNT increases by 60% (ΔF/F =6) upon exposure to 100 μM of DA, which translates to ΔF/F = 0.3 at peak physiological dopamine concentrations that follow burst neuronal firing events (~1 μM).^21,28,29^ Motivated by the goal of producing an *in vivo* compatible neuromodulator nanosensor for a broader dynamic range of physiological relevance, we synthesized a (GT)_N_ based ssDNA-SWNT library for N = 4, 6, 7, 8, 12, 15, 19, 22, 26, and 30 with a previously described protocol.^30^ Near infrared fluorescence and absorption spectroscopy confirm that all sequences from N = 4 to N = 30 produced stable DNA-SWNT suspensions, as evidenced by sharply defined spectral line shapes corresponding to known SWNT electronic transitions (Figure S1). We then measured each (GT)_N_-SWNT nanosensor response to 100 μM dopamine. Consistent with previous results, dopamine addition increases SWNT fluorescence for all sequences (Figure 1). However, there exists a strong length-dependent trend in nanosensor response, for which the previously reported (GT)_15_-SWNT nanosensor represents an apparent minimum (ΔF/F_0_ = 0.45), and (GT)6-SWNT a maximum (ΔF/F_0_ = 24) (Figure 1a, b). ‘Short’ (GT)_N_ polymers (N = 4, 6, 7, 8) yield ΔF/F_0_ = 14, 24, 17, and 10 in response to 100 μM dopamine, respectively, for the (9,4) SWNT chirality. Conversely, ‘long’ (GT)_N_ polymers (N = 12, 15, 19, 22, 26, 30), yield lower ΔF/F_0_ = 0.45, 0.4, 0.5, 0.6,0.4, and 1.5 responses to 100 μM dopamine concentration, respectively (Figure 1b). We identify low baseline fluorescence, F_o_, for ‘short’ (GT)4-8-SWNT complexes as the reason for the large ΔF/F0 values of these constructs (Figure 1b, inset). We also noted that the (GT)_6_-SWNT construct shows increased selectivity towards a new neuromodulator target, norepinephrine, with ΔF/F_0_ = 35 sensitivity (Figure 1c).

**Figure 1.**
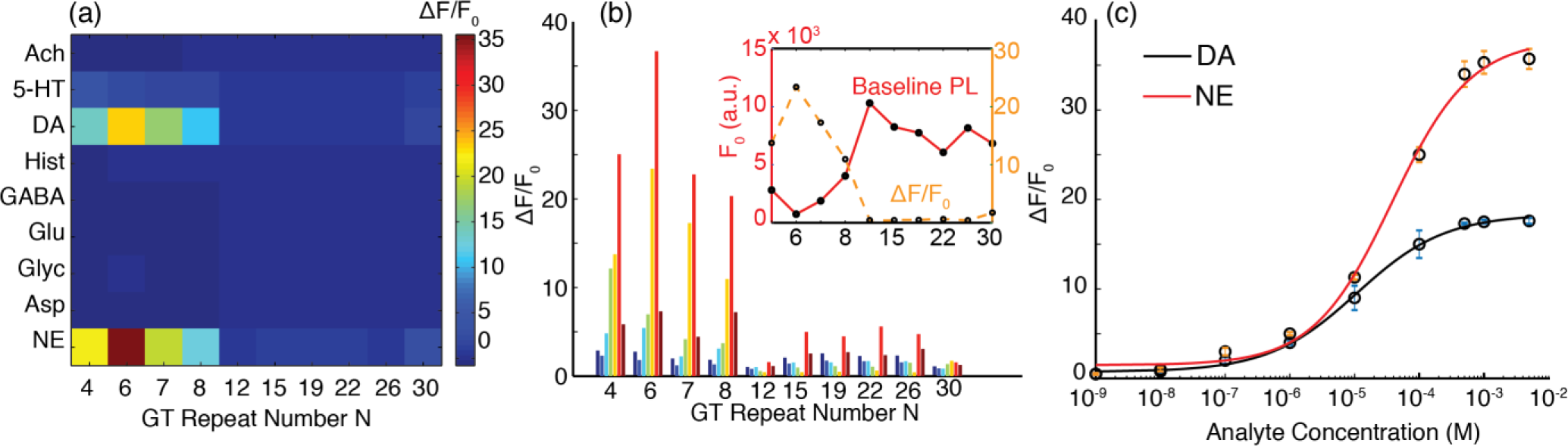
Nanosensor response and selectivity for neuromodulators dopamine and norepinephrine as a function of polymer length. **(a)** Heat map of neurotransmitter screen against (GT)_N_ polymer sequences. Analyte abbreviations: Ach = acetylcholine, 5-HT = serotonin, DA = dopamine, Hist = histamine, GABA = γ-aminobutyric acid, Glu = glutamate, Gly = glycine, Asp = aspartic acid, NE = norepinephrine **(b)** ΔF/F_0_ of each sequence, for each SWNT chirality (8,3) dark blue, (6,5) blue, (7,5) cyan, (10,2) green, (9,4) and (7,6) yellow, (8,6) and (12,1) red, (10,3) and (10,5) maroon. Inset: Baseline fluorescence intensity of (GT)_N_ suspensions at the (9,4) chirality (red) and change in fluorescence intensity after addition of 100 μM of dopamine (orange). **(c)** (GT)_6_-SWNT nanosensor response curve for norepinephrine (red) and dopamine (black). Error bars are standard deviation from n = 3 independent measurements. Experimental data (circles) were fit with Hill equation (solid line).

Our experimental results thus identify polymer length as a key modulator of SWNT fluorescence quantum yield, which can be exploited for maximizing nanosensor sensitivity and improving selectivity for neuromodulators. We further identify the (GT)_6_-SWNT complex as the most suitable nanosensor for imaging both dopamine and norepinephrine, with ΔF/F_0_ = 24 and 35 toward 100 μM analyte concentrations, respectively. DNA-SWNT absorption spectra remain invariant to the addition of dopamine (Figure S1), further suggesting that quantum yield increases drive the increase in nanosensor fluorescence. We next validated the utility of (GT)_6_-SWNT to image dopamine and norepinephrine for *in vivo* relevant concentrations. Concentration-dependent fluorescence response curves for (GT)_6_-SWNT show fluorescence modulations lie within an optimal dynamic range for *in vivo* imaging of neuromodulation (100 nM to 1 μM) (Figure 1c).^29,31,32^ At basal dopaminergic and noradrenergic neuronal activity corresponding to at-rest conditions, we observe that the (GT)_6_-SWNT construct exhibited ΔF/F_0_ values on the order of 1 (100%). At burst firing neuronal activity level typically arising from response to salient events, ΔF/F_0_ values on the order of 5 (500%) can be obtained (Figure 1c). Equally importantly, the high sensitivity imparts the (GT)_6_-SWNT construct an inherently enhanced selectivity for neuromodulators dopamine and norepinephrine over other potentially competing and ubiquitous neurotransmitters, such as glutamate (Glu), acetylcholine (Ach) and γ-aminobutyric acid (GABA) (Figure 1a). We fit our concentration-dependent experimental data points to the Hill equation and determined the dissociation constants (*K_d_*) to be 35 for norepinephrine and 10 μM for dopamine (Figure 1 c).

The molecular selectivity and sensitivity towards catecholamine neuromodulators appears to be highly dependent on nucleobase chemistry. We found that two poly-C sequences, C_30_-SWNT and C_12_-SWNT remain largely non-responsive when exposed to either analyte, consistent with previous studies that show that poly-C ssDNA sequences bind strongly and stably to SWNT (Figure S2).^33^ Other 12-mer sequences, including (GA)_6_, (ATTT)_3_, and (TAT)_4_, similarly exhibit highly diminished sensitivity to both dopamine and norepinephrine (Figure S2). The structure of SWNT surface adsorbed ssDNA is sensitive to charge screening by counter ions^34^ and recent reports have shown that solution ionic strength plays a role in setting the baseline fluorescence (“brightness”) of ssDNA-SWNT constructs.^35^ To rule out ionic strength effects, we tested the response of (GT)_6_-SWNT to both analytes at solution ionic strengths that varied over two orders of magnitude. We found that the turn-on response remained largely insensitive to ionic strength (Figure S2). We also tested the (GT)_6_-SWNT nanosensor response to both analytes at low (pH=4), neutral (pH=7) and high (pH=10) conditions. The fluorescence response is observed at all pH conditions, with best responses observed under physiological pH conditions. Additional experimental evidence was accumulated to substantiate the robustness of the (GT)_6_-SWNT nanosensor. Time-dependent fluorescence (Figure S3) and absorbance (Figure S4) measurements acquired over the course of 7 days confirm polymer-SWNT stability for all values of N except for N=4).

Taken together, these results suggest that the (GT)_6_-SWNT construct can serve as a dopamine and norepinephrine nanosensor with the dynamic range, binding kinetics, and robustness compatible with *in vivo* utility.

## Solvatochromic Shifting Reveals Dopamine and Norepinephrine-Specific Molecular Recognition

We performed surfactant displacement experiments to gain further insight into how analytes modulate the quantum yield of (GT)_N_ functionalized SWNT constructs. Recent work has shown that when added to DNA-SWNT suspensions, surfactants such as sodium cholate (SC) adsorb to exposed SWNT surface, displace adsorbed ssDNA, and alter the SWNT’s surface dielectric properties, thereby causing a solvatochromic shift in exciton optical transition energies (Figure 2a, 2b).^36,37,38^ As expected, addition of SC to (GT)_N_-SWNT induce solvatochromic shifts in (GT)_N_-SWNT fluorescence center wavelengths (Figure 2b). All constructs showed characteristic SC-induced blue-shifting of center wavelengths corresponding to all SWNT chiralities in the sample. We next repeated SC displacement experiments for all (GT)_N_-SWNT suspensions pre-incubated in 10 μM of dopamine. Surprisingly, addition of dopamine to (GT)_N_-SWNT suspensions before addition of SC either reduces or eliminates the SC-induced shifting in exciton optical transitions, suggesting that the surfactant is unable to displace the surface adsorbed ssDNA in the presence of dopamine (Figure 2c, Figure S5c, S5d). We propose that the stabilization of (GT)_N_ polymers on SWNT arises from a selective interaction between the dopamine analyte and dopamine-specific recognition pockets in the (GT)_N_-SWNT conjugate, and that dopamine trapped in binding pockets enhance fluorescence by interacting with both the adsorbed polymer and the SWNT. We posit that as a result of these interactions, polymer-mediated binding of analytes selectively enhances the fluorescence quantum yield of ssDNA-SWNT nanosensors, as we further explore using experimental and computational approaches below.

**Figure 2.**
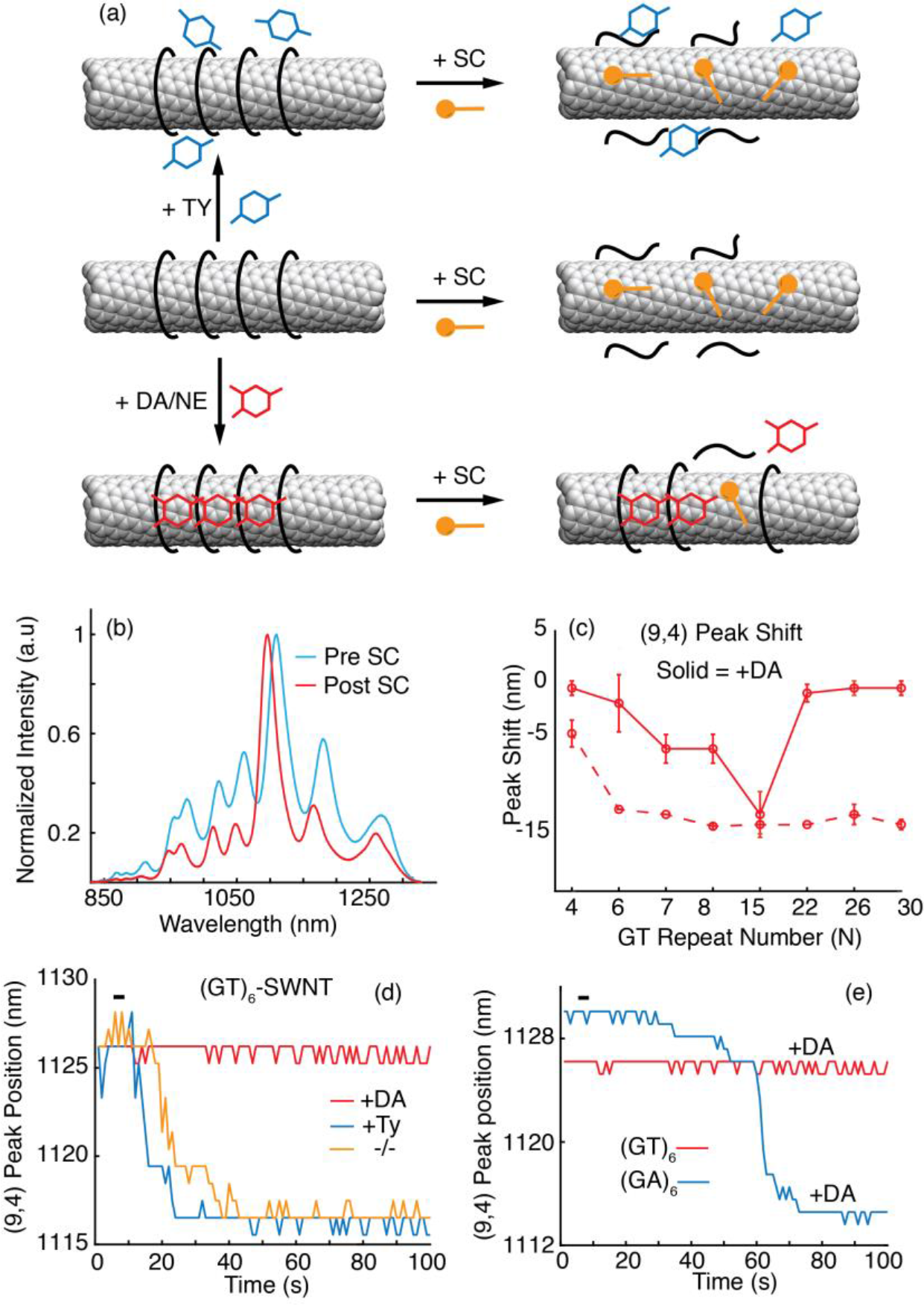
Solvatochromic shifts reveal neuromodulator-specific molecular interactions with nanosensors dependent on ssDNA sequence and length. **(a)** Middle row: sodium cholate (SC) binds to exposed SWNT surfaces and displaces bound (GT)_N_ polymers. Bottom row: Nanosensor incubation in dopamine (DA) or norepinephrine (NE) stabilizes ssDNA polymers on the SWNT surface, disallowing the SC from accessing the SWNT surface. Top row: Incubation in *p*-tyramine (TY) does not stabilize surface adsorbed ssDNA against displacement by SC **(b)** 1% wt.% SC induces a solvatochromic shift in SWNT fluorescence. The shift for the (GT)_8_-SWNT conjugate is presented here as an example. **(c)** Fluorescence peak shift corresponding to the (9,4) SWNT chirality (~1130 nm) upon exposure to 1 wt.% SC without (dash trace) and with (solid trace) incubation in 10 μM of DA. Error bars are standard deviation from n = 3 measurements. Negative peak shits correspond to blue shifting of the peak in the emission spectrum, as shown in (b). **(d)** Time-resolved fluorescence measurements of (GT)_6_-SWNT incubated in 10 μM of DA (red trace), 10 μM of *p*-tyramine (TY) (blue trace), and incubated in neither (orange trace). Upon addition of 0.25 wt % SC indicated by the black bar, peak shift in the dopamine incubated corona is eliminated. **(e)** SC induced solvatochromic peak shift in (GA)_6_-SWNT incubated in 10 μM of dopamine suggests (GA)_6_ exhibits short lived stability on SWNT following dopamine incubation.

To probe the selectivity of dopamine-induced nanosensor stabilization, we conducted time-resolved SC shift experiments with (GT)_6_-SWNT construct in which *p*-tyramine is added to the suspension before addition of SC. Tyramine, a molecular analogue of dopamine differing by only one hydroxyl group, does not modulate the fluorescence of (GT)_6_-SWNT (Figure S6). We reasoned that the recognition of dopamine and norepinephrine is mediated by unique recognition sites in the (GT)_6_-SWNT corona, and that tyramine’s inability to modulate SWNT fluorescence is a consequence of its inability to bind these recognition sites. With this hypothesis, the efficacy of SC in displacing surface adsorbed (GT)_6_ ssDNA and resulting solvatochromic shift should be unaffected by tyramine. Our results do indeed show that 10 μM tyramine, unlike dopamine, does not attenuate the SC induced peak shifts (Figure 2d, Figure S5a), suggesting that tyramine is unable to bind to and stabilize surface adsorbed ssDNA strands.

Our results further indicate that the stability imparted to the SWNT-ssDNA corona phase by the binding of dopamine and norepinephrine is related to the analyte-induced fluorescence modulation specific to the GT base sequence. A (GA)_6_-SWNT construct, in contrast to (GT)_6_-SWNT, exhibits low modulation in fluorescence upon addition of either dopamine or norepinephrine (Figure S2). We also incubated the (GA)_6_-SWNT suspension in dopamine to measure SC induced peak shifts. We observed that dopamine tentatively stabilizes (GA)_6_-SWNT corona (Figure 2e). However, the dopamine-induced stability of (GA)_6_-SWNT is short lived, with distinctive solvatochromic peak shifting occurring with a 60 second delay from SC addition. Another 12-mer sequence, C_12_, similarly exhibits SC-induced solvatochromic shifting despite the presence of dopamine (Figure S5b). These results suggests that SC-induced solvatochromic peak shifting is a function of both the dopamine-bound fraction of recognition sites in the SWNT-polymer corona, and also the intrinsic binding affinity between the polymer sequence and SWNT surface.^37^ Furthermore, we find that both dopamine and norepinephrine modulate the Raman G-band region of (GT)_6_-SWNT between 1500 and 1550 cm^−1^, whereas para-tyramine does not. The effect of Raman G-band broadening by dopamine and norepinephrine, but not tyramine, is maintained regardless of the subsequent addition of SC, suggesting the molecular recognition of dopamine and norepinephrine analytes disallows SC adsorption (Figure S7), and further corroborating our hypotheses.

We probed whether the surface density of the (GT)_N_ polymer on the SWNT surface can tune the density of molecular recognitions sites available to analyte. We varied polymer surface packing of the (GT)_6_-SWNT construct, with different mass proportions of SWNT (mS) to (GT)_6_ DNA polymers (mD). The resulting (GT)_6_-SWNT conjugates thus have variable surface-adsorbed polymer density (Figure S8, Methods) with nominal mS/mD mass ratios of 2, 5, and 10, representing a spectrum from ‘high’ to ‘low’ (GT)_6_ polymer surface density. The resulting fluorescence intensity from equimolar SWNT aliquots shows a clear trend whereby the highest polymer surface densities (mS/mD = 2) exhibit the lowest baseline fluorescence (Figure S8). Addition of 10 μM of dopamine enhances the SWNT fluorescence of all three samples; however, the ΔF/F_0_ nanosensor response is highest for the SWNT sample with the highest surface coverage (Figure S8). These results reveal that (i) the degree of baseline fluorescence quenching of SWNT by adsorbed (GT)_6_ is directly proportional to the polymer surface density; (ii) the higher the polymer surface coverage, the higher the number of dopamine binding pockets; and (iii) dopamine enhances SWNT quantum yield in proportion to the density of bound recognition sites.

## Multiscale Simulations of (GT)_N_ Adsorbed on (9,4) SWNT

We performed multiscale simulations of (GT)_(N=6,15)_-(9,4)-SWNT complexes to disclose mechanisms responsible for a strongly quenched baseline fluorescence and a large nanosensor response to neuromodulators observed in (GT)_6_-SWNT constructs, in contrast to (GT)_15_-SWNT. First, we equilibrated both (GT)_6_-SWNT and (GT)_15_-SWNT systems with atomistic molecular dynamics (MD) simulations. The (GT)15 polymer, which was initially helically wrapped around the SWNT consistent with previous work,^39–43^ remained in a helical conformation during a 200 ns MD simulation (Figure 3a). On the contrary, the (GT)_6_ polymer rearranges from its initial helical conformation into a ring-like conformation in each of the five independent 200 ns trajectories performed. We further examined adsorption of multiple (GT)_6_ on the (9,4) SWNT in a 250 ns long simulation. We observe helix-to-ring transitions in all (GT)_6_ polymers (Figure 3b). The ring conformations of neighboring ssDNAs become highly ordered, as seen from the distinct sharp peaks positioned at approximately equal intervals of ~0.25 nm in the radial distribution function of DNA phosphate groups (Figure S9).

**Figure 3.**
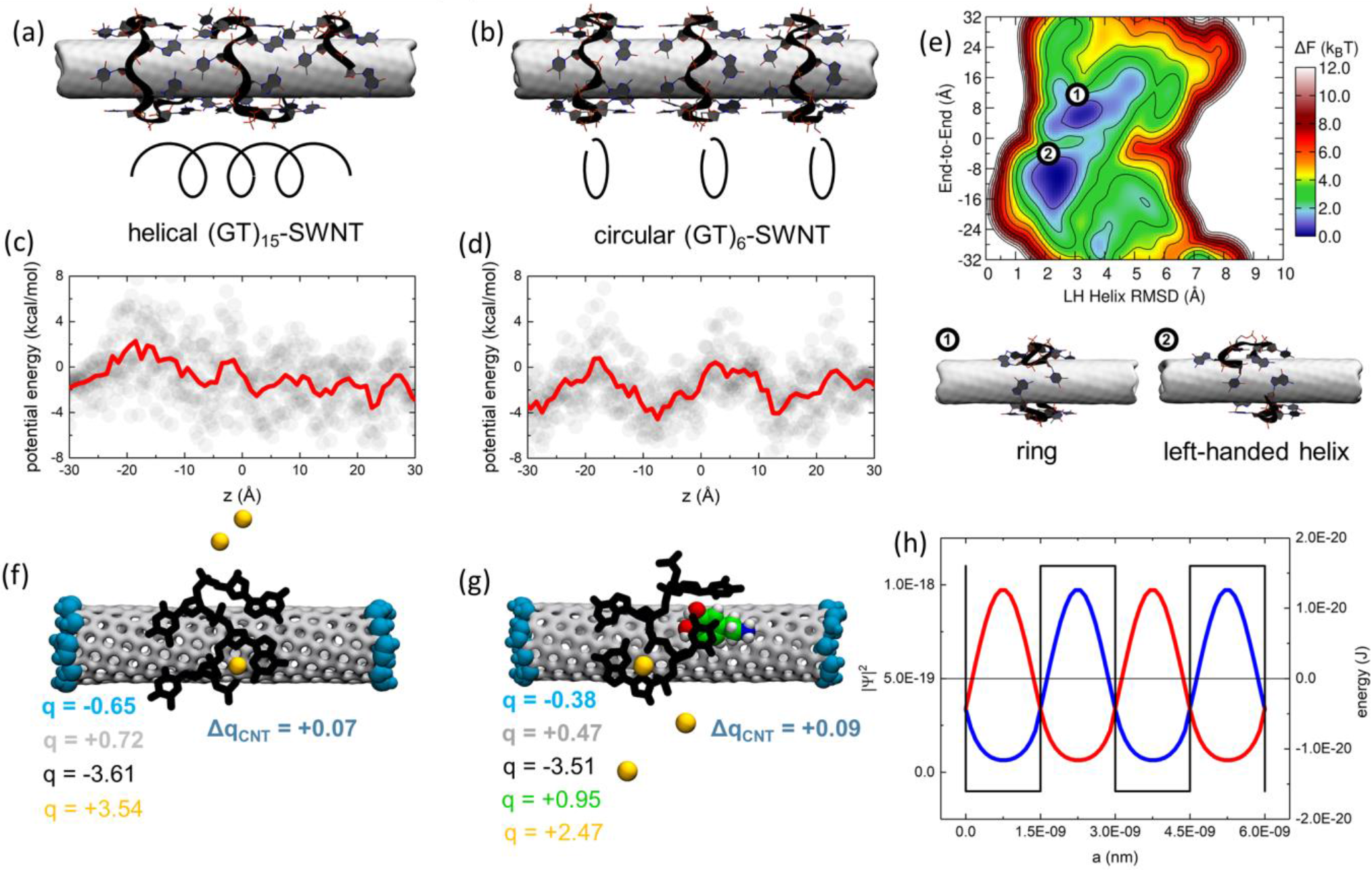
Computational modeling of ssDNA-SWNT nanosensor complexes. **(a)** A representative conformation of (GT)*15*-SWNT. SWNT is shown as a gray surface, (GT)*15* and its backbone are shown in licorice and black ribbon representations, and ssDNA atoms are shown in gray (C), red (O), blue (N) and orange (P). **(b)** A representative conformation of (GT)*6*-SWNT, containing three (GT)*6* polymers. The color scheme is the same as in panel a. **(c)** Electrostatic potential energy profile at SWNT surface in the (GT)*15*-SWNT system as a function of SWNT axial length. The profile is averaged over 2 ns and over the radial SWNT dimension, and includes the effects of the complete SWNT environment present in MD simulations (ssDNA, water and ions). **(d)** Electrostatic potential energy profile at the SWNT surface for the (GT)*6*-SWNT system plotted as a function of SWNT axial length. (e) Free energy landscape of (GT)_6_-SWNT at 300 K. The structures corresponding to two free energy minima are labeled by indices 1 and 2. **(f)** Net charges of molecular fragments in the (GT)_2_-SWNT system, evaluated in quantum mechanical calculations. **(g)** Net charges of molecular fragments in the (GT)_2_-SWNT system with an adsorbed dopamine molecule, evaluated in quantum mechanical calculations. The color scheme in panels f and g: black (DNA), silver (non-terminal SWNT atoms), blue surface (terminal-CH groups capping the SWNT), yellow (sodium ions), green, blue, red and white spheres (C, **N**, O and H dopamine atoms). **(h)** Electron (red) and hole (blue) probability densities in a Kronig-Penney potential (Methods). Probability density values are labeled on the left axis, and the values associated with the potential energy well are labeled on the right axis.

To confirm that the ring-like conformation is a favorable adsorbed state of (GT)_6_ on the (9,4) SWNT, we calculated a free energy landscape of this ssDNA on the (9,4)-SWNT surface at room temperature (T = 300 K) (Figure 3e), using replica exchange molecular dynamics (Methods).^42^ The landscape reveals two distinct stable conformations for (GT)_6_, a left-handed helix and a non-helical ring-like conformation, corresponding to free energy minima at (x,y) = (2.5 Å, −10 Å) and (3.2 Å, 6 Å), respectively, where x represents the root mean square deviations (RMSD) of the DNA structure with respect to the representative left-handed DNA helix, and y represents the distance along the long SWNT axis of two selected atoms of the 3’-and 5’-end DNA nucleotides. These two conformations have approximately the same free energies, and as such they should both be dominant. Moreover, because free energy barrier between them is only ~1.2 kcal/mol, frequent conversions between the two conformations is likely at room temperature for single or sparsely adsorbed polymers. However, in experimental suspensions, SWNT surface is likely to be covered by multiple ssDNA polymers. In that case, the ring-like ssDNA conformations are likely to be more prevalent than the helical conformations due to steric hindrance as they provide better ssDNA surface packing. The ring-like ssDNA conformation is likely enhanced by the fact that the (GT)_6_ contour length matches the circumference of the (9,4) SWNT, affording ordered self-assembly of the oligonucleotides on the SWNT surface.

Since the charged (GT)_6_ and (GT)_15_ polymers have different conformations on the (9,4) SWNT, we reasoned that they should create electrostatic potentials of different topologies close to the SWNT surface. To investigate this phenomenon, we calculated the average electrostatic potential at the SWNT surface generated by all molecules in the system (ssDNA, water, and ions, including the Na^+^ cations adsorbed over long timescales within ssDNA pockets) (Figures S10 and Figure S11). (GT)_15_ creates regions of negative and positive electrostatic potential under the polymer as a ‘footprint’, which extend ~4 nm in contiguous length and roughly follow the ssDNA helical pattern (Figure S12a-b). Negative potential pockets are primarily beneath guanine nucleotides, while positive pockets occur beneath thymine nucleotides. When averaged over the radial SWNT dimension, as shown in Figure 3c, the electrostatic potential profile at the SWNT surface under (GT)_15_ is roughly constant across the entire helix, with random fluctuations. The electrostatic potential around SWNT with adsorbed (GT)_6_ rings also follows the polymer (Figure S12c-d), which results in distinct ring-like regions of alternating positive and negative potentials along the SWNT axis, where each pocket is ~1.5 nm long. In contrast to (GT)_15_-SWNT, when averaged over the radial SWNT dimension (Figure 3d), these electrostatic potentials exhibit large periodic oscillations across multiple rings. Therefore, from the perspective of excitons confined in SWNT’s quasi-1D structure, the periodic electrostatic potentials created by the (GT)_6_ rings effectively form a superlattice (Figure 3d).

Next, QMMD calculations were performed to better understand exciton luminescence in the (GT)_6_-(9,4)-SWNT conjugates (Figure S13). SWNT is polarized by the presence of charged DNA, with overall partial positive charges on SWNT surface covered with ssDNA and partial negative charges at the SWNT ends (Figure 3f). However, we observed only a small charge transfer between ssDNA and SWNT (Figure 3f, Table S1). Dopamine adsorption on the DNA-wrapped SWNT decreased SWNT polarization (Figure 3g and Table S2). However, this effect is only local, and the low molarity of adsorbed dopamine molecules is unlikely to effectively alter the polarizability of a large (GT)_6_-SWNT complex (Figure S14, Figure S15). However, adsorption of dopamine molecules is capable of locally perturbing the periodic electrostatic potential, which will have an effect on SWNT photoluminescence.

These MD and QMMD results provide insight into possible relaxation pathways of excitons in the (GT)_N_-SWNT complexes with and without adsorbed dopamine molecules, and we propose the following mechanisms for the strong turn-on response of (GT)_6_-SWNT nanosensors: (i) SWNT polarization induced by the adsorption of multiple (GT)_6_ polymers can give rise to a dominant non-radiative exciton relaxation mechanism because effective doping activates phonon-assisted relaxation channels for SWNT excitons.^44^ As shown above, the adsorption of dopamine molecules can slightly reduce SWNT polarization but is unlikely to block this non-radiative exciton relaxation mechanism. (ii) Radiative exciton relaxation in SWNT is further attenuated by the presence of closely-spaced periodic potentials of multiple (GT)_6_ strands (Figure 3d). In positive and negative regions of this potential, the electron and hole wave function components tend to avoid each other (Figure 3h, Figure S16, Figure S17), which results in a significant cancelation of their overlap integral present in the oscillator strength.^45^ Therefore, in a (GT)_6_-(9,4)-SWNT complex with high SWNT coverage, radiative transitions of excitons are expected to be significantly suppressed, in agreement with the low baseline photoluminescence observed in experiments. However, in the presence of adsorbed dopamine molecules, the cancellation of the overlap integral can be disturbed (disordered superlattice) and radiative transitions can be active simultaneously with the non-radiative transitions, giving rise to a large-magnitude fluorescent turn-on nanosensor.

## Conclusions

In this work, we report (GT)_6_-SWNT as a strong turn-on optical nanosensor for the neuromodulators dopamine and norepinephrine with a dynamic range compatible for applications to the study of *in vivo* neurophysiology. We investigated the photophysical and molecular underpinnings of the strong and selective turn-on response. We conclude that SWNT-ssDNA nanosensors with selective fluorescence modulation towards an analyte exhibit selectivity through specific binding interactions involving the SWNT, the adsorbed polymer, and the analyte. We further show that the magnitude of a nanosensor turn-on response can be tuned by varying polymer contour length and adsorption surface density. We used multi-scale computational approaches to rationalize our experimental findings. Molecular dynamics simulations reveal that self-assembly of (GT)_6_ ssDNA on the SWNT surface produce highly ordered ring structures, which effectively form a superlattice from the perspective of a 1-D confined SWNT exciton. The resultant periodic potential promotes a dominant non-radiative relaxation of excitons and dim baseline SWNT fluorescence, which can be selectively enhanced by an analyte via perturbation of the superlattice that promotes competitive radiative relaxation. These insights and results have important implications for new nanosensor discovery for other biomolecular analytes of interest, as well as for orthogonal fields of research hinging on ssDNA-SWNT interactions such as SWNT purification by chiral index.

## Acknowledgments

We would like to thank Dr. Michael Ross and the Peidong Yang lab at University of California, Berkeley, Department of Chemistry for help with Raman measurements. We acknowledge startup funding from the University of Texas at El Paso (to A. A. A., L.V.), the NSF Division of Materials Research, Grant # 1506886 (to P.K.), Burroughs Wellcome Fund Career Award at the Scientific Interface (CASI) (M.P.L), the Simons Foundation (M.P.L), a Stanley Fahn PDF Junior Faculty Grant with Award # PF-JFA-1760 (M.P.L), a Beckman Foundation Young Investigator Award (M.P.L), and a DARPA Young Investigator Award (M.P.L). M.P.L. is a Chan Zuckerberg Biohub investigator. A.G.B. is supported by an NSF Graduate Research Fellowship and an NIH F99/K00 award from NINDS.

